# Mapping Activity-Dependent Quasi-Stationary States of Mitochondrial Membranes with Graphene-Induced Energy Transfer Imaging

**DOI:** 10.1101/2021.06.15.448547

**Authors:** Sufi Oasim Raja, Alexey I. Chizhik, Christoph F. Schmidt, Jörg Enderlein, Arindam Ghosh

**Affiliations:** Department of Physics and Soft Matter Center, Duke University, Durham, NC 27708, United States; Third Institute of Physics—Biophysics, University of Göttingen, Göttingen, Germany; Cluster of Excellence “Multiscale Bioimaging: from Molecular Machines to Networks of Excitable Cells” (MBExC), Georg August University, 37077 Göttingen, Germany

**Keywords:** graphene-induced energy transfer, mitochondrial membranes, IM-OM distance, hyper-osmotic shock

## Abstract

Graphene-induced energy transfer (GIET) was recently introduced for the precise localization of fluorescent molecules along the optical axis of a microscope. GIET is based on near-field energy transfer from an optically excited fluorophore to a single sheet of graphene. As a proof-of-concept, we demonstrated its potential by determining the distance between the two leaflets of supported lipid bilayers (SLBs) with sub-nanometer accuracy. Here, we use GIET imaging for three-dimensional reconstruction of the mitochondrial membrane architecture. We map two quasi-stationary states of the inner and outer mitochondrial membranes before and during adenosine tri-phosphate (ATP) synthesis. We trigger the ATP synthesis state in vitro by activating mitochondria with precursor molecules. Our results demonstrate that the inner membrane (IM) approaches the outer membrane (OM) while the outer membrane (OM) does not show a measurable change in average axial position upon activation. As a result, the inter-membrane space (IM-OM distance) is reduced by ∼2 nm upon activation of the mitochondria. This direct experimental observation of the subtle dynamics of mitochondrial membranes and the change in inter-membrane distance induced by ATP synthesis is relevant for our understanding of the physical functioning of mitochondria.

---

**T**he development of super-resolution fluorescence microscopy (SRFM) has enabled the imaging of sub-cellular organelles with unprecedented detail. ^1–4^ The most widely used class of SRFM is single-molecule localization microscopy (SMLM), comprising methods such as photo-activated localization microscopy (PALM), ^5^ stochastic optical reconstruction microscopy (STORM), ^6^ fluorescent PALM (fPALM), ^7^ direct STORM (dSTORM), ^8^ or point accumulation for imaging in nanoscale topography (PAINT). ^9,10^ These methods can determine lateral positions of single fluorescent emitters with a resolution of few to tens of nanometers. For localizing the position of single molecules also along the optical axis of a microscope (and thus enabling 3D SMLM), several schemes such as astigmatic imaging, biplane imaging, or point spread function engineering have been developed and successfully implemented. ^11–18^ Nevertheless, in most cases the achievable axial resolution remains three to five times worse than the lateral resolution. Only interferometric techniques such as iPALM ^19^ or isoSTED ^20^ offer an isotropic nanometric resolution along all three dimensions, albeit at the price of elevated technical complexity. Metal-induced energy transfer (MIET) was developed as a simple alternative for axial localization of fluorophores with nanometer accuracy using a conventional fluorescence lifetime imaging microscope (FLIM). MIET relies on an electrodynamic nearfield mediated energy transfer from an optically excited fluorescent emitter (donor) to plasmons in a thin planar metal film (acceptor). ^21–24^ This energy transfer is strongly distance dependent, leading to a characteristic modulation of a fluorophore’s excited-state lifetime as a function of its distance to the metal, within a total range of up to ∼250 nm from the metal surface. Using an appropriate theoretical model, a measured lifetime can then be translated into an axial distance value which forms the basis of MIET (axial) super-resolution imaging.

Recently, we have shown that by replacing the metal layer with a single sheet of graphene, the resolution of MIET can be enhanced nearly tenfold, thus enabling sub-nanometer optical localization of fluorophores at reasonable photon budgets. We demonstrated the potential of graphene-induced energy transfer (GIET) by determining the distance between the two leaflets of supported lipid bilayers (SLBs). ^25^ Subsequently, GIET imaging was used for investigating the nanoscopic conformational organization and dynamics of membrane-anchored Rab7-like GTPase Ypt. ^26^ In the current work, we utilize GIET imaging for quantifying subtle dynamic changes in mitochondrial membrane organization by mapping the distance of the inner and outer membranes of mitochondria from a graphene sheet before and during ATP synthesis.

Mitochondria, the power (ATP) plants of cells, play a vital role in cellular function, ^27,28^ and mitochondrial dysfunction is involved in many neuro-degenerative diseases. ^29^ Structurally, a mitochondrion is a double-membrane organelle. The outer membrane (OM) that encloses the organelle has a composition that is similar to the composition of the cellular plasma membrane. The inner membrane (IM) is highly invaginated and forms structures known as cristae, which serve as the central functional units of mitochondria. Cristae are the sites of biochemical reactions related to ATP synthesis, and previous reports suggest that the density of invaginations in cristae is directly correlated with the functionality of a mitochondrion. ^30^ Live-cell STED nanoscopy has shown that mitochondrial cristae are highly dynamic structures. ^31^ However, it is not known whether mitochondrial membranes undergo dynamic changes (in particular the inter-membrane spacing) when switching from the resting state to ATP synthesis.

GIET imaging offers a unique opportunity to map the axial position of mitochondrial membranes with nanometer resolution along the axial direction. For this purpose, we isolated mitochondria from HEK 293 cells (see supporting information for details). We designed two different types of fluorophore-tagged mitochondria. In type I, we stained the inner-membrane (IM) with the commercially available dye mitotracker deep red (MTDR). This probe has been previously utilized for STORM imaging of the inner membrane of mitochondria. ^1^ In type II mitochondria, the outer membrane was fluorescently labeled by tagging the protein complex TOMM20 with the fluorescent protein mCerulean3 (see supporting information for details of fluorophore labeling). GIET measurements were then performed as follows: a GIET substrate was prepared by evaporating a layer of SiO_2_ on top of a graphene-coated glass coverslip (see supporting information for details). Isolated activatable mitochondria were then immobilized on top of the GIET substrate. To facilitate attachment of the mitochondrial membrane via electrostatic interactions, the surface was coated with a uniform layer of positively charged poly-L-lysine (PLL). The complete scheme of the experiment is shown in figure 1A (see supporting information for details). In order to map the axial distance of the membranes from the substrate, FLIM images of the mitochondria were recorded (see figure S1 in supporting information). An exemplary FLIM image of each type of mitochondria is presented in figure 1B. The measured fluorescence lifetime values were translated into axial distances using a theoretically calculated GIET curve, i.e. the functional dependence of fluorescence lifetime on distance (see figure S2 in supporting information). This calculation requires knowledge of the fluorescence quantum yield and the free-space lifetime of the fluorophores. These values were obtained *a priori* in independent experiments, for both MTDR and mCerulean3. Precise quantum yield values of MTDR and mCerulean3 were determined through a nanocavity-based method, ^32^ and the obtained values are *ϕ* = 0.30 ± 0.02 and *ϕ* = 0.19 ± 0.01, respectively. Using time-correlated single-photon counting (TCSPC), the free-space lifetime of MTDR and mCerulean3 were determined to be *τ*_0_ = 1.94 ± 0.08 ns and *τ*_0_ = 3.71 ± 0.10 ns, respectively.

**Figure 1.**
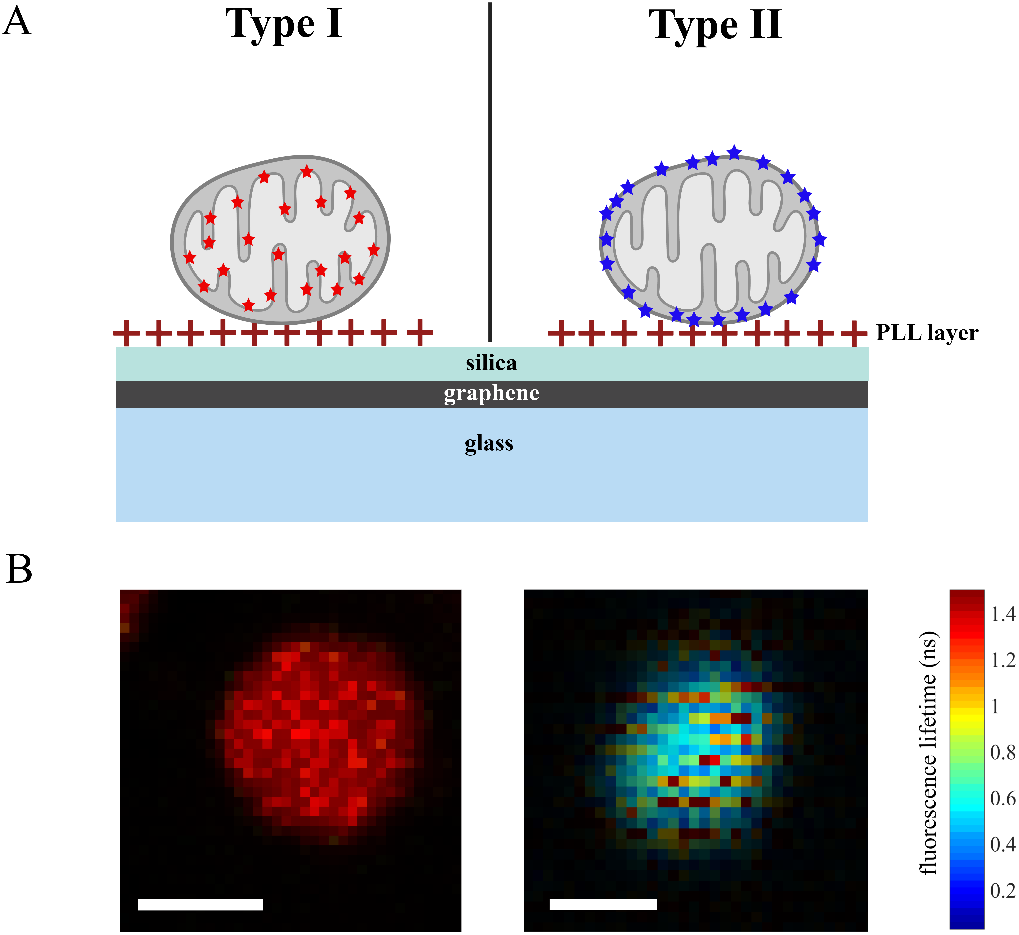
A Geometry of the experiment: The schematic shows isolated mitochondria that are functionally immobilized on a GIET substrate. The substrate consists of a silica layer of 5 nm thickness evaporated on top of a single sheet of graphene deposited on a cover slip. A layer of poly-L-lysine (PLL) (shown as **+**) is put on top of the silica to enable better attachment of mitochondria via electrostatic interaction. Type I (left) represents mitochondria where the inner membrane is labeled with mitotracker deep red (MTDR) (red stars). Type II (right) represent mitochondria where the protein complex TOMM-20 residing in the outer membrane was labeled with mCerulean-3 (blue stars). **B** Exemplary FLIM images of a type I (left) and a type II (right) mitochondrion, functionally immobilized on a GIET substrate. Scale bar 1 µm.

Isolated mitochondria contain all the machineries to start respiration (oxygen consumption) and ATP synthesis through oxidative phosphorylation. Therefore, isolated mitochondria can be made active and ATP synthesis can be triggered by using suitable precursor molecules. ^33^ We mapped the average membrane-surface distance of mitochondria in two different states. We refer to these two states of mitochondria as ‘resting’ and ‘active’ states. In the active state, mitochondria are induced to synthesize ATP by immersing them into a buffer containing precursors of ATP synthesis, while the resting state is observed before such activation. The activation buffer contains pyruvate, malate, and adenosine di-phosphate (ADP). The composition of this activation buffer has been optimized according to mitochondrial metabolism and function as reported in. ^34^ We recorded FLIM images of isolated mitochondria stained with MTDR and converted the measured fluorescence lifetime values into IM-substrate distance values (see figure S2 and supporting information for further details). We monitored the IM-substrate distance of the same mitochondria in their resting state and after activation with the respiration buffer (see figure 2A). These measurements show that the IM-substrate distance decreases upon activation. Furthermore, we observed a change in the shape of mitochondria upon activation. We also recorded FLIM images of a set of mitochondria (*N* = 35) in their resting state which show an average IM-substrate distance of 12.42 *±* 0.71 nm (see Figure 2B). Similar measurements for mitochondria after activation show an average IM-substrate distance of 10.40 *±* 0.70 nm (Figure 2B). Thus, activation of mitochondria shifts the inner membrane by ∼2 nm towards the substrate.

**Figure 2.**
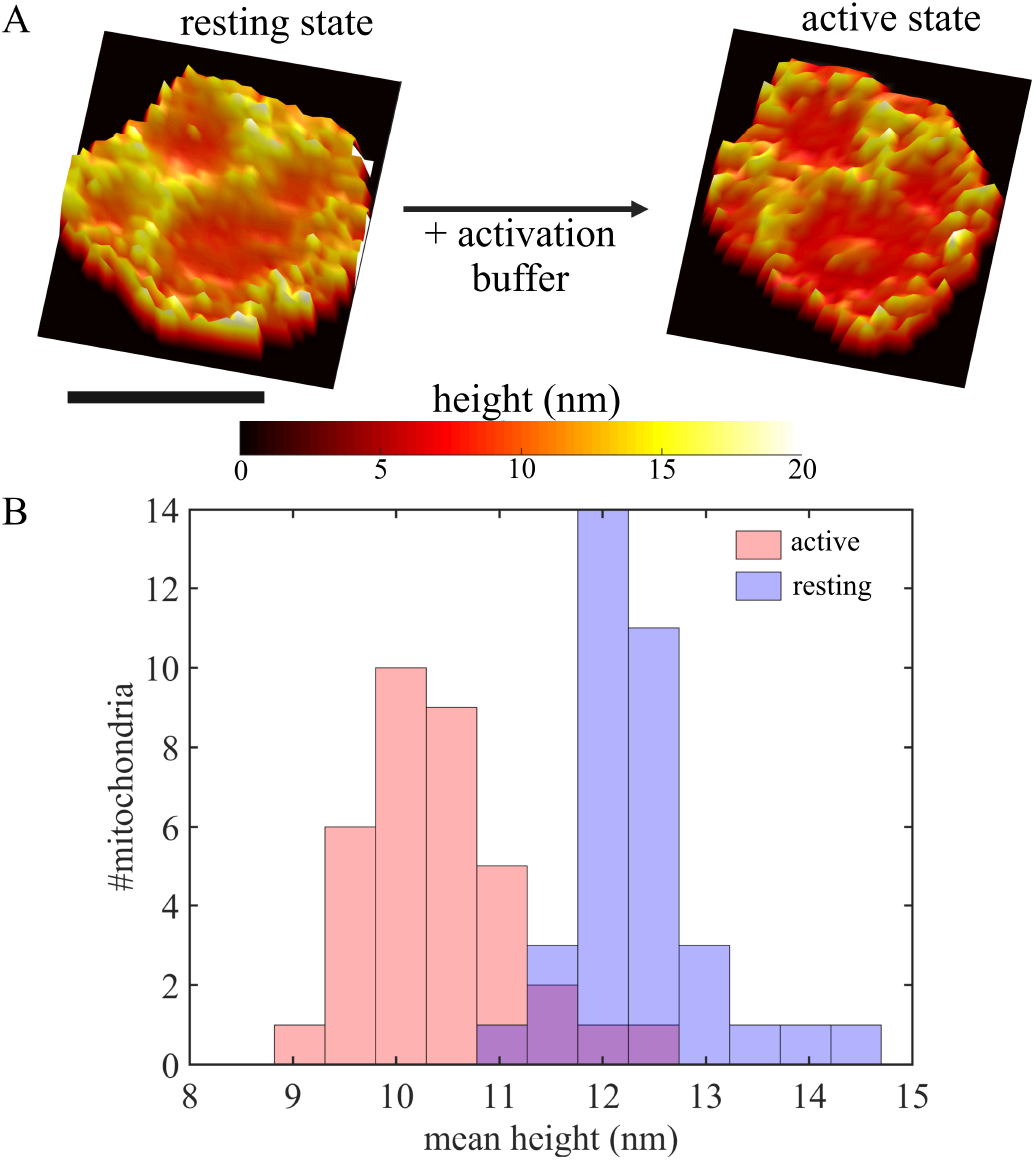
The inner membrane approaches the surface with activation: GIET imaging of a type I mitochondrion (inner membrane labeled with MTDR). **A** Three-dimensional reconstruction of the inner membrane of a representative mitochondrion imaged in the resting state (left) and after activation (right). Scale bar 1 µm. **B** Average distance of mitochondrial inner membrane from the substrate as obtained from imaging a set of 35 mitochondria in their resting state, and another set of 35 mitochondria after activation. For mitochondria imaged in their resting state, we obtain an average distance of the inner membrane to the surface of 12.42 *±* 0.71 nm. For mitochondria imaged after activation, the average distance of the inner membrane from the surface was found to be 10.40 *±* 0.70 nm.

Next, we focused on type II mitochondria for which the outer membrane was fluorescently labeled. Using GIET imaging, we did not observe any significant change in the OM-substrate distance of these mitochondria from resting to active states. Figure 3A shows three-dimensional reconstructions of the outer membrane of a set of mitochondria before and after activation. We repeated these measurements on two different batches of type II mitochondria, where one batch (*N* = 33) was imaged in its resting state and the other after activation. The average OM-substrate distance was found to be 1.85 ± 0.41 nm and 1.85 ± 0.37 nm at resting and activated states, respectively. Using these values, we can estimate the IM-OM distance or thickness of the inter-membrane space in the resting and active states. For mitochondria in their resting state, this distance was found to be 10.57 ± 0.58 nm, while it reduces to 8.55 ± 0.59 nm in the active state. Previous studies using electron microscopy (EM) on fixed mitochondria ^35,36^ reported an inter-membrane distance in the order of 6-12 nm. However, EM studies are not able to discern the average IM-OM distance of functional mitochondria during and before ATP synthesis. Here, GIET imaging with its extraordinary axial resolution is a unique technique for mapping the quasi-stationary states of mitochondrial membranes before and during ATP synthesis.

**Figure 3.**
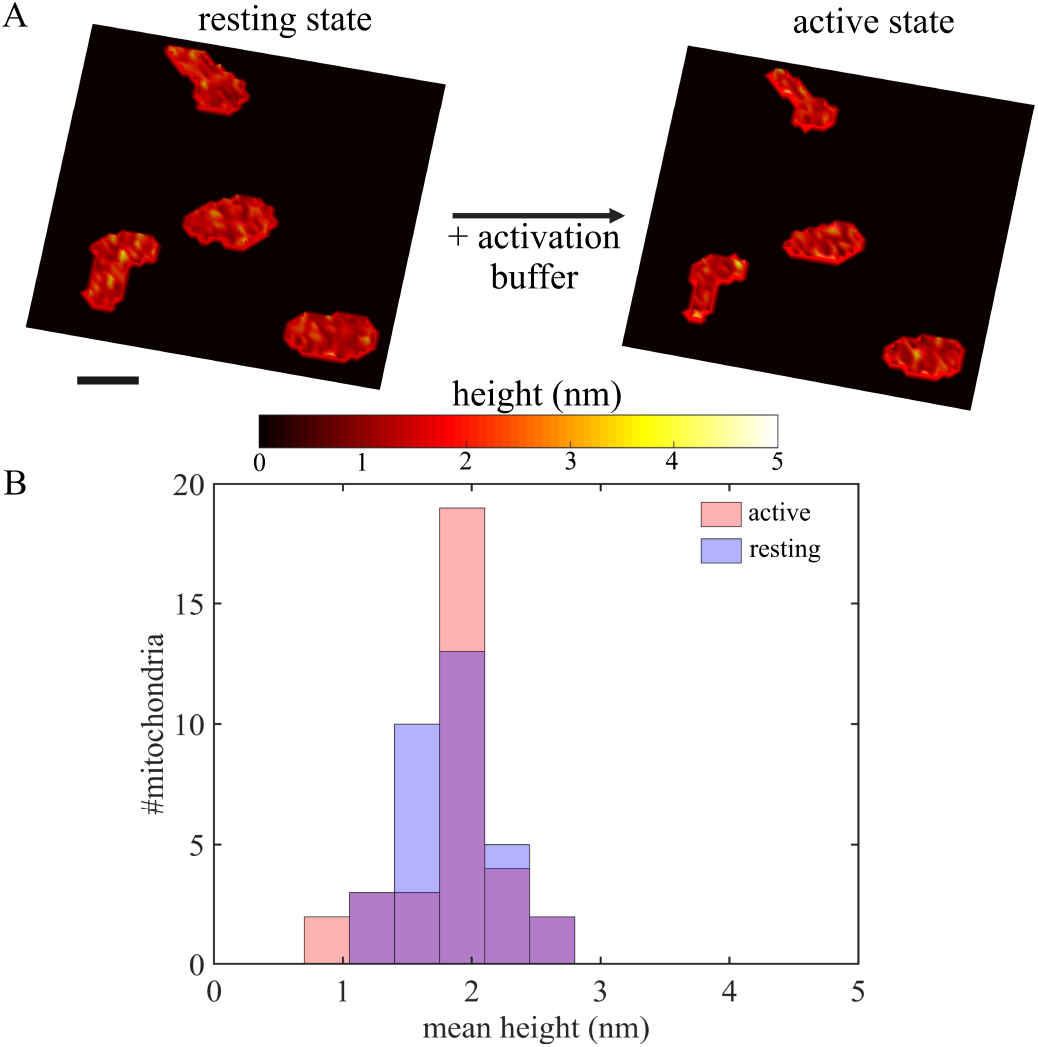
The outer membrane maintains same distance from the surface during activation: GIET imaging of type II (TOMM-20 tagged with mCerulean3 in outer membrane) mitochondria. **A** Three-dimensional reconstruction of the outer membrane of four mitochondria imaged in the resting state (left), and the same mi-tochondria imaged after activation (right). Scale bar 1 µm. **B** Average distance of mitochondrial outer membrane from the substrate as obtained from imaging a set of 33 mitochondria in their resting state, and another set of 33 mitochondria after activation. For mitochondria in their resting state, we obtain an average distance of the outer membrane of 1.85 ± 0.41 nm. For mitochondria after activation, the average distance of the outer membrane from the surface was found to be 1.85 *±* 0.37 nm.

To further validate our results, we applied hyper-osmotic shocks to the mitochondria after activation. We used 0.5 M trehalose, a non-reacting complex sugar for this purpose. Confocal micrographs, as presented in figure 4A and plots from line scans in figure 4B, demonstrate the shrinkage of the inner volume of mitochondria after hyper-osmotic shock. Figure 4C present FLIM images on a GIET substrate of type I mitochondria at three different stages - in resting, activated, and post hyper-osmotic shock state (from left to right). We observed a reduction in fluorescence lifetime of MTDR in active as compared to resting mitochondria. This observation is in excellent agreement with our GIET imaging results as presented earlier. Next, the same set of mitochondria was exposed to hyper-osmotic shock which resulted in an increase of fluorescence life-time values of MTDR accompanied by a shrinkage of the inner volume (right panel of figure 4C). Time-correlated single-photon counting (TCSPC) histograms and average fluorescence lifetimes of a set of type I mitochondria after hyper-osmotic shock are shown in figure S3. De-quenching of fluorescence lifetime after hyper-osmotic shock reflects the fact that the inner membrane is moving further away from the GIET substrate as compared to its distance in the active state. Since fluorescence lifetime values after hyper-osmotic shock approach the free-space lifetime of MTDR, a height reconstruction from FLIM images is no longer possible in this case. To cross-check possible effects of activation on hyper-osmotic shock, we applied a hyper-osmotic shock also to type I mitochondria in their resting state. We observed a similar shift of fluorescence lifetimes towards longer values, which excludes any possible effect of the respiration buffer on hyper-osmotic shock (see figure S4 and experimental details in supporting information).

**Figure 4.**
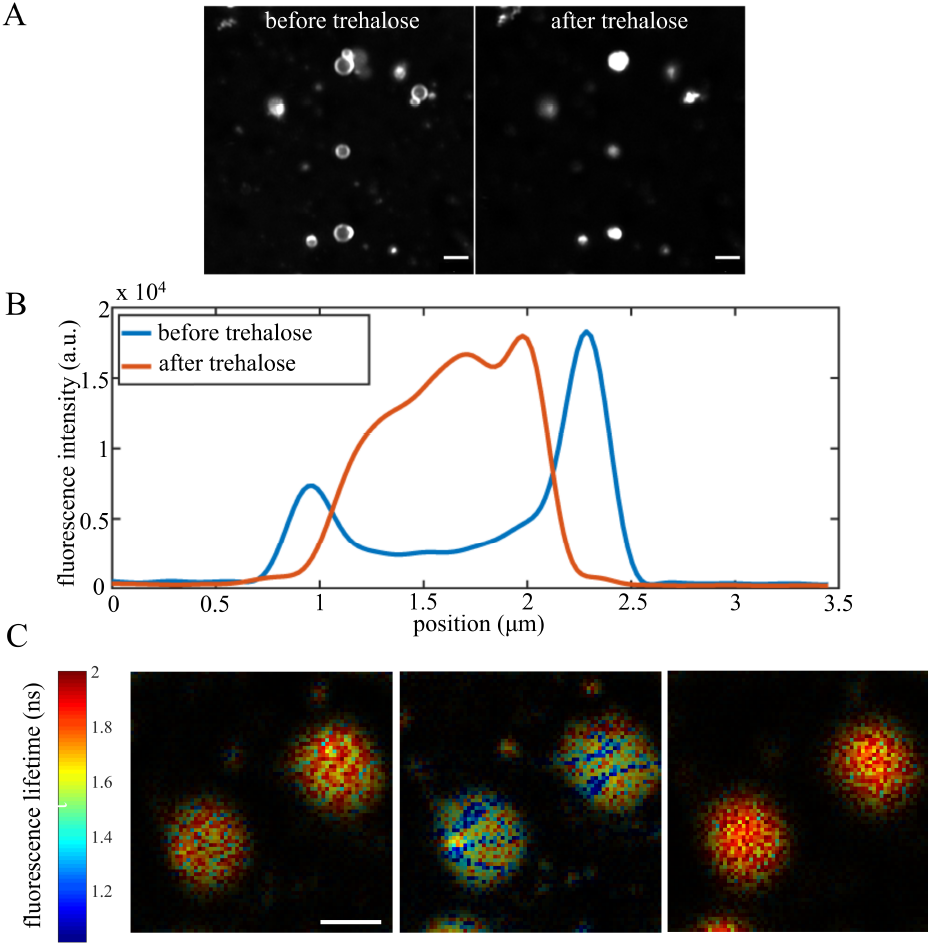
Hyper-osmotic shock induced by addition of trehalose. **A** Confocal micrograph of MTDR-stained isolated mitochondria before and after trehalose-induced hyper-osmotic shock. Scale bar 2 *µ*m. See video S1 in supporting information. **B** Line scans along the mitochondria clearly indicate shrinkage of the inner volume of mitochondria. **C** FLIM image of two type I mitochondria on a GIET substrate in their resting state (left). The same mitochondria imaged after treatment (middle panel) with activation buffer, exhibiting a reduction in fluorescence lifetime as compared to the same sample in the resting state. Again, the right panel shows a FLIM image of the same mitochondria after addition of 0.5 M trehalose. As compared to the middle panel, longer fluorescence lifetime values close to the free-space lifetime of MTDR are observed upon hyper-osmotic shock with trehalose. Scale bar 1 µm.

To summarize, we utilized GIET imaging for measuring the distance between the IM and OM of mitochondria in their resting and active states. Cristae are dynamic structures ^37^ that become more numerous and ordered upon increase in mitochondrial activity.^38^ It is plausible that cristae reorganization, involving an increase in IM surface area, leads to a change in inter-membrane spacing if the organelles maintain inner-volume homeostasis, which is typically a crucial and highly regulated characteristic of cells. While mitochondrial volume regulation itself plays a very important role in cell function, ^39^ one can hypothesize that the activity-dependent change in inter-membrane spacing is an efficient way of regulating the flux of metabolites. To conclude, GIET imaging with its exceptional axial resolution is an emerging technique that allows for elucidating subtle structural changes in membrane organization (and potentially protein-membrane interaction) in sub-cellular organelles.

## ASSOCIATED CONTENT

### Supporting Information

The Supporting Information is available free of charge. GIET substrate preparation, isolation of mitochondria, fluorescent labeling of mitochondrial membranes, sample preparation, FLIM imaging on isolated mitochondria, fluorescence lifetime data evaluation, calculation of GIET calibration curves, conversion of lifetimes into axial distances, hyper-osmotic shock experiments, height maps and figures S1 - S5 are provided.

## Acknowledgements

S.O.R acknowledges light microscopy core facility (LMCF) of Duke University. S.O.R and C.F.S. thank Start-up funding from Duke University. J.E. is grateful for support by the Deutsche Forschungsgemeinschaft (DFG, German Research Foundation) under Germany’s Excellence Strategy, EXC 2067/1-390729940. Financial support from German Research Foundation (DFG, SFB 860, project A06) is gratefully acknowledged by A.G.

