## Supporting figures S1, S2, S3, S4, S5. for "Mapping Activity-Dependent Quasi-Stationary States of Mitochondrial Membranes with Graphene-Induced Energy Transfer Imaging"

#### GIET substrate preparation

Plasma cleaned glass coverslips (24 mm × 24 mm, thickness 170 $\mu$m) coated with a graphene monolayer (0.34 nm thickness) were purchased from Graphene Supermarket, New York, USA. A $SiO_2$ spacer of thickness 5 nm was evaporated on top of this coverslip using an electron beam source (Univex 350, Leybold) under high vacuum conditions ($10 \times 10^{-6}$ mbar). Slowest rate of deposition was maintained (1 Å $s^{-1}$) to ensure maximal homogeneity. The spacer thickness was continuously monitored during evaporation with an oscillating quartz unit.

#### Isolation of mitochondria

We used a commercially available kit (Thermo Fisher Scientific, cat. no. 89874) for isolating mitochondria from cultured cells. We followed the guidelines provided by the manufacturer. Briefly, HEK-293 cells (ATCC® CRL-1573) were lysed by lysis buffer (comes with the kit) and mitochondria were isolated through differential centrifugation. Isolated mitochondria were stored using a special storage buffer as reported earlier[1] to keep the mitochondria functionally intact for long time. Mitochondria were snap-frozen in liquid nitrogen and stored at -80 °C.

#### Fluorescent labeling of mitochondrial membranes

HEK-293 cells were transfected by mCerulean3 tagged TOMM-20 (Addgene, 55449) to stain the outer membrane of mitochondria and inner membrane was stained by MitoTracker Deep Red FM dye, MTDR (Thermo Fisher Scientific, M22426). Cells were transfected by a liposome-based method using Lipofectamine transfection reagent (Thermo Fisher Scientific, L3000001). Figure S1 shows representative confocal FLIM images of stained isolated mitochondria.

#### Sample preparation

10 µL frozen mitochondria were slowly thawed in ice and mixed with 20 µL of storage buffer. Diluted mitochondria suspension was added on top of poly-l-lysine coated GIET surface. Plates were incubated at 4 °C for half an hour so that mitochondria can stick well to the surface. Next, 70 $\mu$l of respiration buffer containing 125 nM of MTDR was added. After 15 min of incubation at room temperature we started lifetime imaging. For activation assay we prepared a master mix as described earlier[2] in respiration buffer containing 1 mM malate (Sigma Aldrich), 5 mM pyruvate (Sigma Aldrich) and 100 µM ADP (Sigma Aldrich).

### FLIM imaging on isolated mitochondria

FLIM imaging was performed on a home-built confocal setup equipped with an objective lens of high numerical aperture (Apo N, 100× oil, 1.49 NA, Olympus Europe). The excitation unit

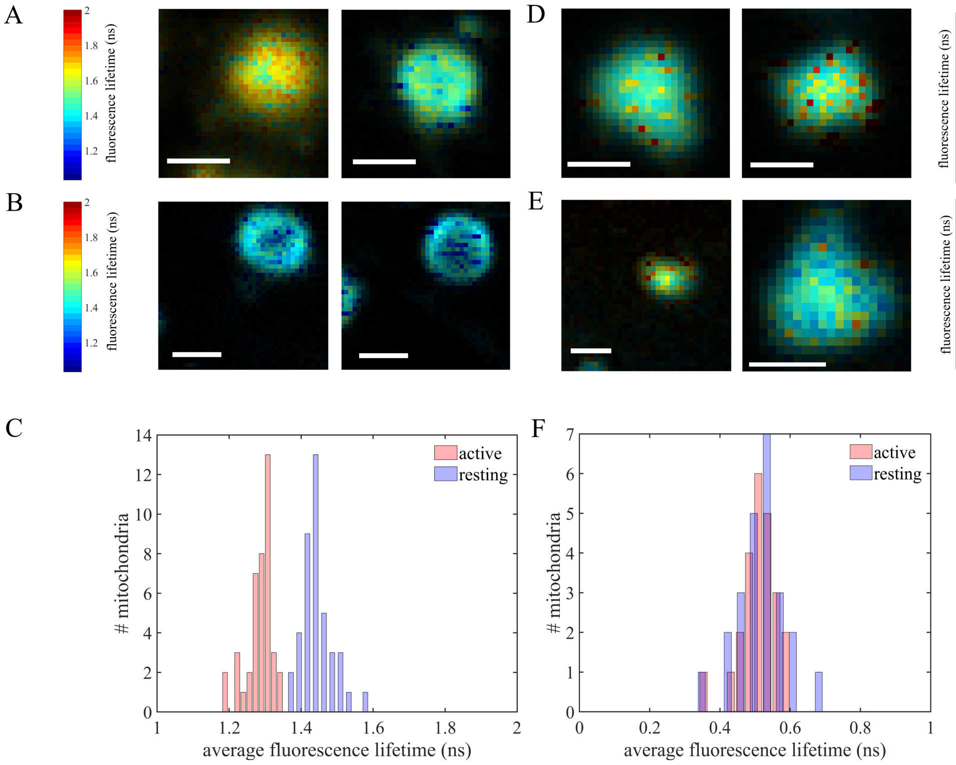


Figure S1: FLIM imaging on isolated mitochondria. **A** FLIM images of two exemplary type I mitochondria in their resting state. **B** FLIM images of two different type I mitochondria in their active state. As can be seen, fluorescence lifetime of MTDR in the inner membrane of mitochondria in active state (**B**) is shorter as compared to resting state (**A**). **C** Bar histogram of fluorescence lifetime values as obtained from two different sets of resting (red) and active (blue) type I mitochondria. **D** FLIM images of two type II mitochondria in their resting state. **E** FLIM images of two different type II mitochondria in their active state. As can be seen, fluorescence lifetime of mCerulean3 in the outer membrane of mitochondria in active state (**E**) doesn't change as compared to resting state (**D**). **F** Bar histogram of fluorescence lifetime values as obtained from two different sets of resting (red) and active (blue) type II mitochondria.

consisted of a pulsed, linearly polarized white light laser (SC400−4−8, Fianium Ltd., pulse width ∼50 ps, repetition rate 80 MHz) equipped with a tunable filter (AOTFnC 400.650-TN, Pegasus Optik GmbH). For our measurements, we used 460 nm and 640 nm wavelengths for fluorescence excitation of mCerulean3 and MTDR respectively. Excitation light was reflected by a non-polarizing beam-splitter towards the objective lens. Back-scattered excitation light was blocked using long-pass filters (EdgeBasic BLP01-488R-25, Semrock for the blue channel, and BLP01-635R, Semrock, for the red channel). Emission light was focused onto the active area of a SPAD-detector (SPCM-AQRH, Excelitas) with an achromatic lens (AC254-030-A-ML, Thorlabs), and data recording was done with a multi-channel picosecond event timer (HydraHarp 400, PicoQuant GmbH). Isolated mitochondria were scanned with a focused laser spot using a piezo nanopositioning stage (P-562.3CD, Physik Instrumente GmbH). Exemplary FLIM images of mitochondria on GIET substrate is illustrated in the main text in figure 1B. We also present FLIM images of both types of mitochondria in their resting and active states in figure S1. Distribution of average fluorescence lifetime values obtained from multiple mitochondria are depicted in figures S1C and S1F.

### Fluorescence lifetime data evaluation

As mentioned previously, acquisition of FLIM images was performed using a multi-channel picosecond event timer (HydraHarp 400, PicoQuant GmbH) in time-tagged, time-resolved (TTTR) mode. It means that for each photon, both the absolute time of the detection ('macro-time') and the time delay between the previous excitation pulse and the detection ('micro-time') are known. We group the micro-times of all photons of a single sample point into a histogram , whose tail was fitted with a multi-exponential function. The lifetimes used in this work are the inverse of the average decay rate of the multi-exponential fit.

### Calculation of GIET calibration curves

GIET calibration curves were calculated as described previously.[3] The geometry of our experiments is depicted in figure 1A in main text. A fluorophore is located at a distance $z_0$ above a GIET substrate. We perform fluorescence excitation and detection through that substrate, from below. We model the electrodynamic coupling of the fluorophore and graphene by treating the fluorescent molecule as an ideal electric dipole emitter and graphene as a layer of matter with specific thickness and (complex-valued) bulk refractive index. Solving Maxwell's equations for such a system then leads to an expression for the emission power, $S(\theta,z_0)$, of the electric dipole emitter as a function of dipole distance $z_0$ and orientation (described by the angle $\theta$ between the dipole axis and the vertical axis).[4–6] The emission

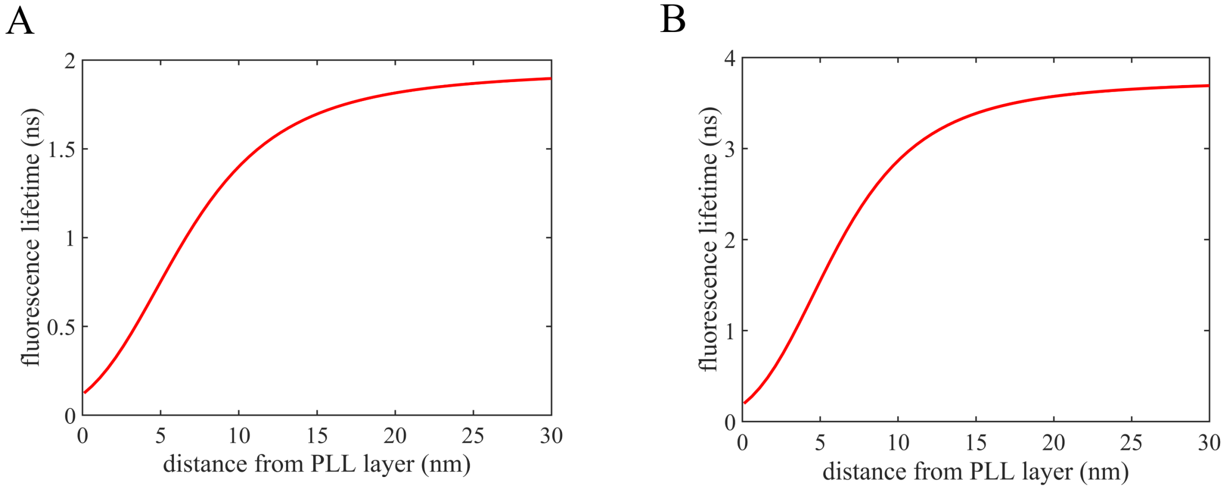


Figure S2: Figure shows calculated GIET model curves for **A** MTDR stained to inner mitochondrial membrane and **B** mCerulean3 tagged to TOMM-20 in outer mitochondrial membrane. For MTDR, measured quantum yield value $\phi = 0.30$ and free-space lifetime $\tau_0 = 1.94$ ns were used with an emission maximum of 665 nm. For mCerulean3, measured quantum yield value $\phi = 0.19$ and free-space lifetime $\tau_0 = 3.71$ ns were used with an emission maximum of 475 nm.

power $S(\theta, z_0)$ itself is inversely proportional to the radiative transition rate of the molecule's excited state to its ground state, and together with the non-radiative rate due, for example, to collisions with surrounding molecules, defines the measurable excited-state (fluorescence) lifetime $\tau_f$:

$$\frac{\tau_f(\theta, z_0)}{\tau_0} = \frac{S_0}{\phi S(\theta, z_0) + (1-\phi)S_0} \tag{1}$$

Here, $\phi$ is the quantum yield (QY), $\tau_0$ is the free-space lifetime in the absence of GIET, $S_0$ is the free-space emission power of an ideal electric dipole emitter. We measured the QY and fre-space lifetimes of the fluorophores tagged to mitochondrial membranes separately as already mentioned in the main text. Figure S2A and S2B shows the calculated relative lifetime

($\tau_f/\tau_0$) as a function of distance $z_0$ for MTDR (staining inner membrane) and mCerulean3 (tagged to TOMM20 in outer membrane) respectively. For these and all subsequent calculations, we adopted a graphene layer thickness of 0.34 nm and a refractive index of $n_{graphene}$ = 2.68 + 1.17$i$ (corresponding to an emission wavelength of 475 nm) and $n_{graphene}$ = 2.76 + 1.40$i$ (corresponding to an emission wavelength of 665 nm).[7] The thickness of the silica layer (refractive index $n_{SiO_2}$ = 1.46) above the graphene was set to 5 nm and an additional 1 nm was considered for the PLL layer. Above it, the refractive index of the membrane was considered to be $n_{graphene}$ = 1.46 as reported earlier for plasma membrane.[8] As can be seen, graphene-induced fluorescence quenching / modulation takes place within the first ∼25 nm. The fluorophores were considered as randomly oriented as it was determined previously for dye molecules staining the cell membrane.[9]

#### Conversion of lifetimes into axial distances

The theoretical background of calculating GIET and MIET curves for converting measured lifetime values into substrate-fluorophore distances is described above and has been published in further detail elsewhere.[3,9] Lifetime values, as obtained from FLIM experiments, can be translated into axial distances using our our MATLAB - based open - source software MIET-GUI. It is available online https://projects.gwdg.de/projects/miet/ repository/raw /MIET_GUI.zip?rev=ZIP. Briefly, one needs to define the sample parameters first by providing thickness and refractive indices of the layers (e.g. graphene, silica), emission wavelength, quantum yield ($\phi$) and free-space lifetime ($\tau_0$). Next, the .mat file containing fitted lifetime values and a separate .mat file containing fluorescence intensity values in the FLIM image has to be uploaded. Check the box indicating 'Visualize height profile' and then click 'Evaluate'. Three dimensional reconstruction of the height image, FLIM image, and fluorescence intensity image are generated as outputs.

### Hyper-osmotic shock experiments

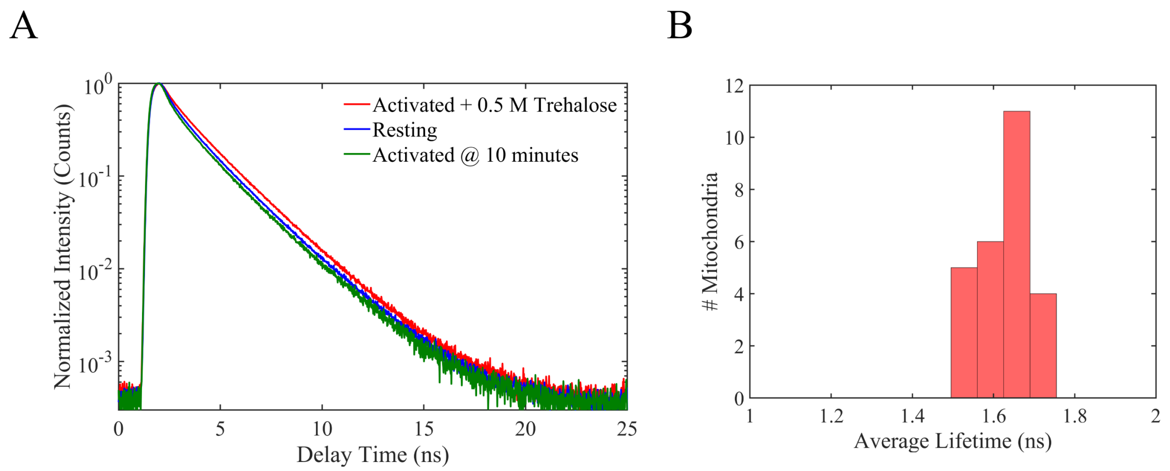


Figure S3: **A** shows TCSPC histograms of MTDR of a set of mitochondria (shown in figure 4C of main text) in resting (blue), activated (green) and post hyper-osmotic shock (red) to activated state of the mitochondria. As can be seen, fluorescence decay becomes shorter from resting to active state and reverts back to longer decay upon treatment with trehalose inducing hyper-osmotic shock. **B** Histogram of average lifetimes of MTDR in trehalose-treated mitochondria in active state as obtained from a set of $N = 25$ mitochondria.

Hyper-osmotic shock was applied to mitochondria in active state using a non-reacting complex sugar trehalose (0.5 M). Figure 4A in the main text demonstrates shrinking of the inner volume of mitochondria upon treatment with trehalose. Consequently, fluorescence lifetime of MTDR in the inner membrane increases near a GIET substrate after hyper-osmotic shock as compared to lifetime values in active state (figure 4C). Figure S3A demonstrates the time-correlated single-photon counting (TCSPC) histograms of those mitochondria shown in figure 4B of main text. Furthermore, histogram of average lifetimes of a set of $N = 25$ mitochondria are presented in figure S3B. As can be seen, the lifetime values approach free-space lifetime upon hyper-osmotic shock. This observation confirm a shrinkage in the inner volume of mitochondria upon hyper-osmotic shock which results in shifting of the fluorophores further away from the GIET substrate. We also verified the effect of trehalose-treatment on mitochondria in resting state to cross-check any effect of activation on hyper-osmotic shock. Figure S4A shows FLIM image of a MTDR-stained mitochondrion in resting state.

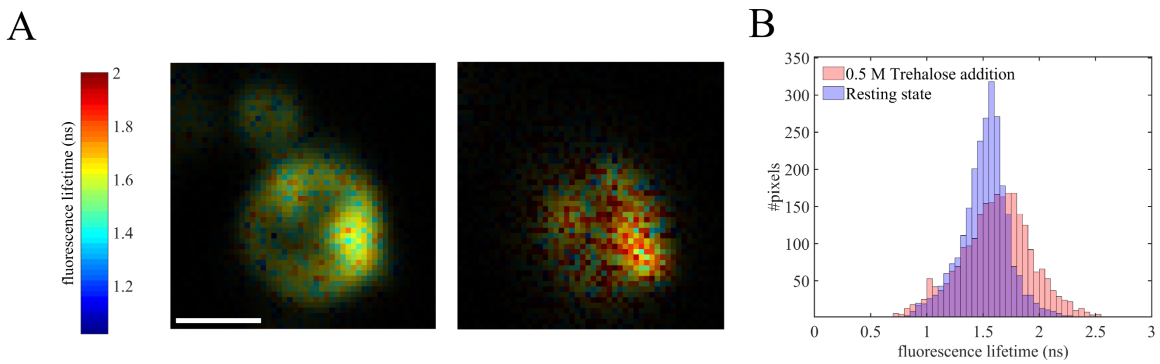


Figure S4: **A** FLIM image of a MTDR-labeled mitochondrion in resting state (left panel) and the same after hyper-osmotic shock. **B** Lifetime distributions as obtained from **A** showing a clear shift of lifetime to longer values upon hyper-osmotic shock. Scale bar 1 $\mu$m

We observe a clear shift in fluorescence lifetime to longer values after hyper-osmotic shock (right panel) as compared to before trehalose treatment in resting state (left panel). The corresponding lifetime distributions are presented in figure S4B. These results eliminate any possible influence of activation on hyper-osmotic shock.

### Height maps

Figure S5A demonstrates three-dimensional height reconstruction of a representative set of type I mitochondria in their resting state. A different set of type I mitochondria were imaged in their active state and their three-dimensional reconstruction of axial distances are shown in figure S5B.

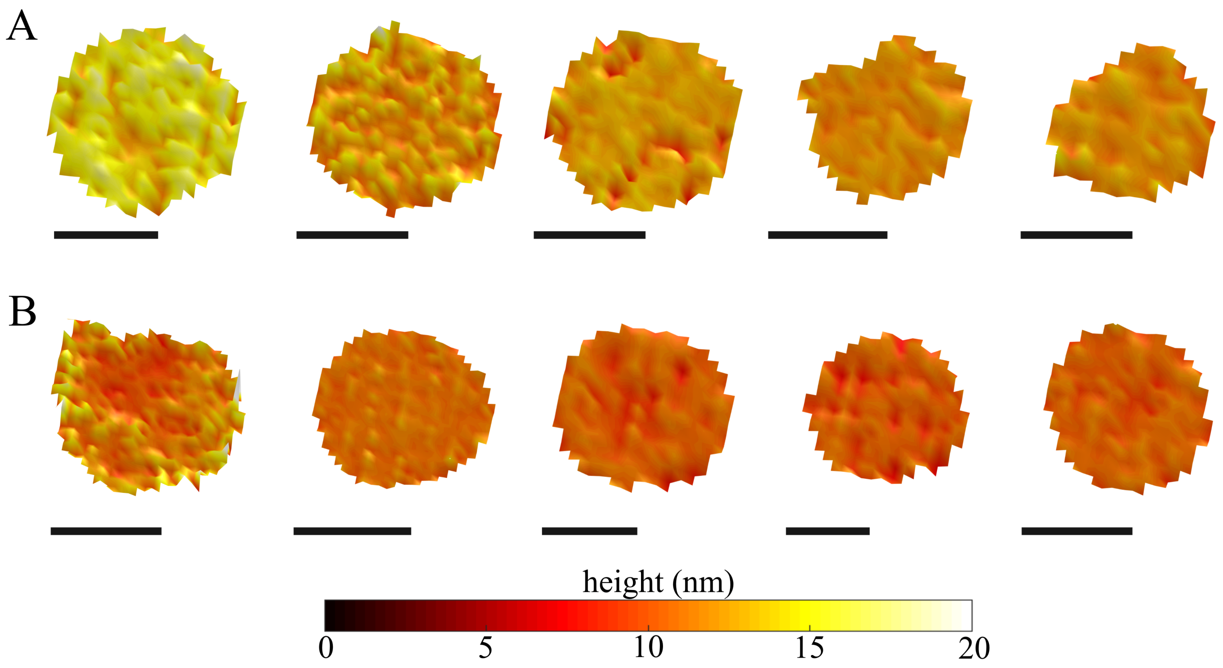


Figure S5: **A** Three-dimensional height maps of five type I mitochondria (IM stained with MTDR) imaged in their resting state. **B** Same for a different set in their active state. All scale bars 1 $\mu$m
